# LoCS-Net: Localizing Convolutional Spiking Neural Network for Fast Visual Place Recognition

**DOI:** 10.1101/2024.03.14.584997

**Authors:** M. Ugur Akcal, Ivan Georgiev Raikov, Ekaterina Gribkova, Anwesa Choudhuri, Ivan Soltesz, Rhanor Gillette, Girish Chowdhary

## Abstract

Visual place recognition (VPR) is the ability to recognize locations in a physical environment based only on visual inputs. It is a challenging task due to perceptual aliasing, viewpoint and appearance variations and complexity of dynamic scenes. Despite promising demonstrations, many state-of-the-art VPR approaches based on artificial neural networks (ANNs) suffer from computational inefficiency. Spiking neural networks (SNNs), on the other hand, implemented on neuromorphic hardware, are reported to have remarkable potential towards more efficient solutions computationally, compared to ANNs. However, the training of the state-of-the-art (SOTA) SNNs for the VPR task is often intractable on large and diverse datasets. To address this, we develop an end-to-end convolutional SNN model for VPR, that leverages back-propagation for tractable training. Rate-based approximations of leaky integrate-and-fire (LIF) neurons are employed during training to enable back-propagation, and the approximation units are replaced with spiking LIF neurons during inference. The proposed method outperforms the SOTA ANNs and SNNs by achieving 78.2% precision at 100% recall on the challenging Nordland dataset, compared with 53% SOTA performance, and exhibits competitive performance on the Oxford RobotCar dataset while being easier to train and faster in both training and inference when compared to other ANN and SNN-based methods.

## 1 Introduction

Visual place recognition (VPR) is the ability to recognize locations in a physical environment based only on visual inputs. It is essential for autonomous navigation of mobile robots, indoor assistive navigation aid, augmented reality, and geolocalization [1, 2, 3, 4, 5, 6]. These applications generally involve complex dynamic scenes, perceptual aliasing, viewpoint and appearance variation, which render VPR extremely challenging.

VPR has been approached via deep learning techniques [7, 8, 9] and various supervised and selfsupervised feature descriptor representations [10, 11, 12]. Despite the promise of such methods, many of them suffer from severe computational inefficiencies [13]. Such limitations significantly reduce the ability for real-world deployment of conventional artificial neural networks (ANNs) on robotic platforms with limited on-board resources [14]. Spiking neural networks (SNNs) offer an alternative with their remarkable potential for computationally efficient operation when they are implemented on neuromorphic hardware [15]. However, previous work on SNN models for VPR has suffered from scalability problems that impede their application to data with a large number of locations. In addition, the majority of the aforementioned methods formulate VPR as an image retrieval task [16], the solution of which aims for the correct association of given query images with a set of reference images. Such formulation requires the employment of a confusion matrix (a.k.a. distance matrix) [17] populated with similarity scores based on the distances between model-specific feature descriptors. A commonly-used similarity metric is the cosine similarity [18], which is reported to be computationally expensive when evaluating high-dimensional feature vectors [19].

These drawbacks have motivated our approach to VPR, described in this paper, in which an SNN model uses an ANN-to-SNN conversion method in order to enable training with backpropagation and yields fast training and inference time. We employ a smooth rate-based approximation [20] of the leaky integrate-and-fire (LIF) neurons [21] during the training. Once the training session is completed the rate-based units are substituted with the spiking LIF neurons and the resulting spiking network is used for inference.

We formulate VPR as a classification task, where the SNN model predicts place labels that uniquely correspond to the locations in a discretized navigation domain. We evaluate our method with the challenging real-world benchmark datasets Nordland [22] and Oxford RobotCar [23, 24].

Our model, the Localizing Convolutional Spiking Neural Network (LoCS-Net), outperforms other state-of-the-art VPR methods based on both ANNs and SNNs on the Nordland dataset [22], while exhibiting competitive performance on the Oxford RobotCar dataset [23, 24].

The main contributions of this work are as follows:

- To the best of our knowledge, LoCS-Net is the first SNN that is trained to perform the VPR task by means of ANN-to-SNN conversion and backpropagation.
- LoCS-Net is an end-to-end SNN solution. Therefore, LoCS-Net doesn’t require further processing of its outputs for recognizing the places. In that sense, LoCS-Net saves all the computation resources that would be spent by traditional VPR algorithms for feature encoding, descriptor matching, computing the similarity scores, and storing a distance matrix.
- We demonstrate that our proposed VPR algorithm yields the fastest inference time among its SNN counterparts. This poses LoCS-Net as a significant step towards deployment of SNN-based VPR systems on robotics platforms for real-time localization.

## 2 Related work

### Task-specific feature descriptors

are the very core of the traditional VPR systems, which can be grouped into two categories: 1) Local descriptors, 2) Global descriptors. Local descriptors may scan the given images in patches of arbitrary size and stride. These patches are then compared to their immediate neighborhood to determine the distinguishing patterns[25]. In general, previous VPR work utilizing the local descriptors [26, 27, 28] employs sparse filters such as [29, 30] that extract so-called key-points. These key-points can be marked by the descriptions generated through the application of methods including SIFT [31], RootSIFT [32], SURF [33], and BRIEF [34]. In this way, the combination of heuristics-based detectors and local descriptors can be used for: A) Representing images, B) Comparing two images with respect to their descriptors to determine how similar they are. In addition, one can concatenate local features with other embeddings [35] while leveraging their robustness against the variations in the robot’s pose. However, local descriptors can be computationally heavier and more sensitive to the illumination changes [36]. Global descriptors [37, 38], on the other hand, do not require a detection phase and directly encode the holistic properties of the input images. Although this might save the global descriptor-based VPR methods [39, 40, 41, 42] some compute time, they are more vulnerable to robot pose changes compared to their local descriptor-based counterparts while being inept at capturing geometric structures[43]. Yet, still the global descriptors are reported to be more effective in the case of varying lighting conditions[44]. Furthermore, there are hybrid approaches [45, 46, 47], which combine strengths of both approaches.

### Conventional ANN-based VPR methods

Deep learning has made key contributions to recent work on VPR. An influential deep-learning-based approach is NetVLAD [48], which is a supervised method for place recognition, based on the Vector of Locally Aggregated Descriptors (VLAD), a technique to construct global image features representations from local feature descriptors. NetVLAD uses a pre-trained feature extraction network, such as AlexNet[49], to extract the local features, and a loss function that aims to minimize the distance between a baseline input and the most similar image (the positive example), while maximizing the distance between baseline input and the most dissimilar image (the negative example). This loss function is also known as the triplet loss function. Several authors have extended NetVLAD in different directions, and NetVLAD-based methods still perform very competitively [47, 50].

### SNNs

have been of interest for various robotics tasks, including not only VPR, but also object detection[51], regression[52], and control of aerial platforms[53] due to their significant potential for computational efficiency[54]. Published VPR methods based on SNNs are relatively recent, compared to other robotics research areas. Among them, [55] is reported to be the first high-performance SNN for VPR. There, the authors propose a feed-forward SNN, where the output neuron activations are filtered through a custom softmax layer. Follow-up work by the same authors [56] introduced a framework where localized spiking neural ensembles are trained to recognize places in particular regions of the environment. They further regularize these networks by removing output from “hyper-active neurons”, which exhibit intense spiking activity when provided input from the regions outside of the ensemble’s expertise. This framework yields a significant improvement over its predecessor while demonstrating either superior or competitive VPR performance compared to the traditional methods. However, training of these SNN approaches do not scale with the increasing volume of training data. In addition, heuristics such as the assignment of neural ensembles to spatial regions, nearest neighbor search in the similarity matrix, and the regularization process further complicate the training process and the computational efficiency of the model. In contrast to these previous SNN-based approaches, we propose an end-to-end solution that is much easier to train and to deploy without requiring heuristic training.

## 3 LoCS-Net Model for Visual Place Recognition

Here, we begin with an overview of the task formulation and the architecture of LoCS-Net in Section 3.1 Section 3.2 formally poses the VPR problem as a classification task. Then, in Section 3.3, we walk through the LoCS-Net pipeline and its key design choices. Moreover, Section 3.3 provides a summary of the ANN-to-SNN conversion paradigm while elaborating on its use for the present work. We defer further design details and the hyper-parameters to the supplementary information.

### 3.1 Overview

Fig. 1 depicts the overall architecture of LoCS-Net. The input to the model is a set of images sampled along a trajectory that traverses a bounded navigation domain. The domain is discretized by means of a uniform grid (orange lines in Fig. 1) and each image is assigned an integer *place* label based on the tile traversed at the time of sampling the image. In this manner, we define the VPR task as a classification problem as discussed in Section 3.2.

**Figure 1.**
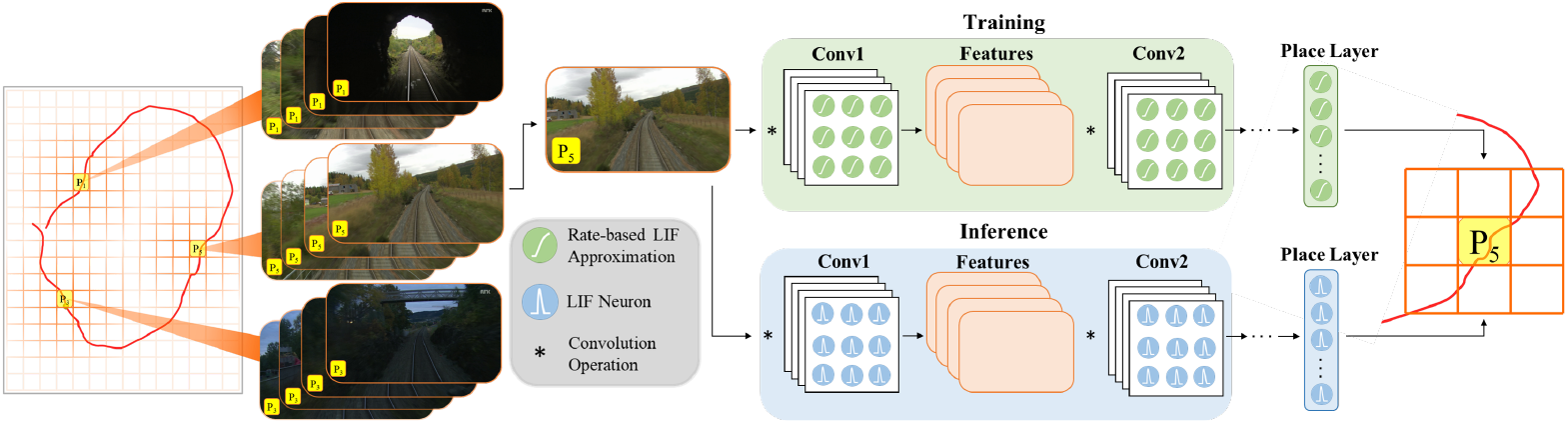
LoCS-Net VPR system. A convolutional network of rate-based LIF neurons [20] is trained over a set of annotated images sampled over a trajectory (the red curve) traversing a finite discretized (orange grid bounded by gray lines) navigation domain. The VPR task is formulated as a classification problem where each tile of the grid (*P*_1_, *P*_2_, *P*_3_,…) corresponds to a distinct location. After training, the LIF approximations are substituted with the spiking LIF neurons [21] for the inference step.

Each layer in the LoCS-Net model consists of Leaky Integrate-and-Fire (LIF) neurons [21]. In order to train the model, these neurons are converted to rate-based approximations of LIF [20]. Rate-based LIF approximations are continuous differentiable representations of the LIF activation function. The LIF activation function describes the time evolution of the neuron’s membrane potential, and it is discontinuous: when the membrane potential reaches a threshold value, it is reset back to a predetermined state. The rate-based approximation is a continuous function that describes the neuron’s firing rate as a function of its input, enabling the use of back-propagation algorithms for training. However, this doesn’t prevent the substitution of the approximate LIF neurons with the original ones for inference after the training is complete. A number of authors have reported successful applications [57, 58, 59] of ANN-to-SNN conversion.

### 3.2 VPR as a classification task

A common practice to approach the VPR task is to pose it as an image retrieval problem where the goal is to compute and store descriptors that would effectively encode both the set of query images and the collection of reference images to match [60]. The encoding process is followed by an image retrieval scheme, which is based on comparing query embeddings (z_q_) to the database of reference descriptors (z_r_) with respect to the customized similarity metrics. Nevertheless, computation of the descriptors is numerically expensive. In contrast, we formulate the VPR task as a classification problem in order to bypass the encoding phase of the images. We designed the LoCS-Net so that it would uniquely map the given input images to the mutually exclusive classes, which are the distinct places, as discussed in Sections 3.1 and 3.3.

Fig. 2 shows how our work formulates VPR differently, as compared to the image retrieval VPR formulation. We first discretize the navigation domain by using a uniform rectangular grid (Fig. 2-b, the orange lines). Here, each tile of the grid defines a distinct place *P*_*i*_, *i* = 1, 2, 3, … . We would like to note that the navigation domain can be any physical environment with points described by spatial coordinates. Although we use a uniform rectangular grid to discretize the top-down view of the domain of interest, our approach is flexible with respect to the definition of places, and permits 3-D as well as 2-D discretization. As one of many ways to generate the training and test data, we sample images over numerous trajectories traversing the discretized navigation domain. Suppose that an image *s ∈* 𝒮 is sampled at the time instant when the camera is in the region represented by tile *P*_5_. Then, this image would be annotated by the place label 5 *∈* 𝒴. Namely, the image *s* belongs to the class represented by the tile *P*_5_. Thus, given a query image, our goal is to train a spiking neural network model that would correctly infer the associated place labels. Hence, we pose the VPR task as an image classification problem in this fashion. We now formally describe the VPR task as a classification problem as follows.

**Figure 2.**
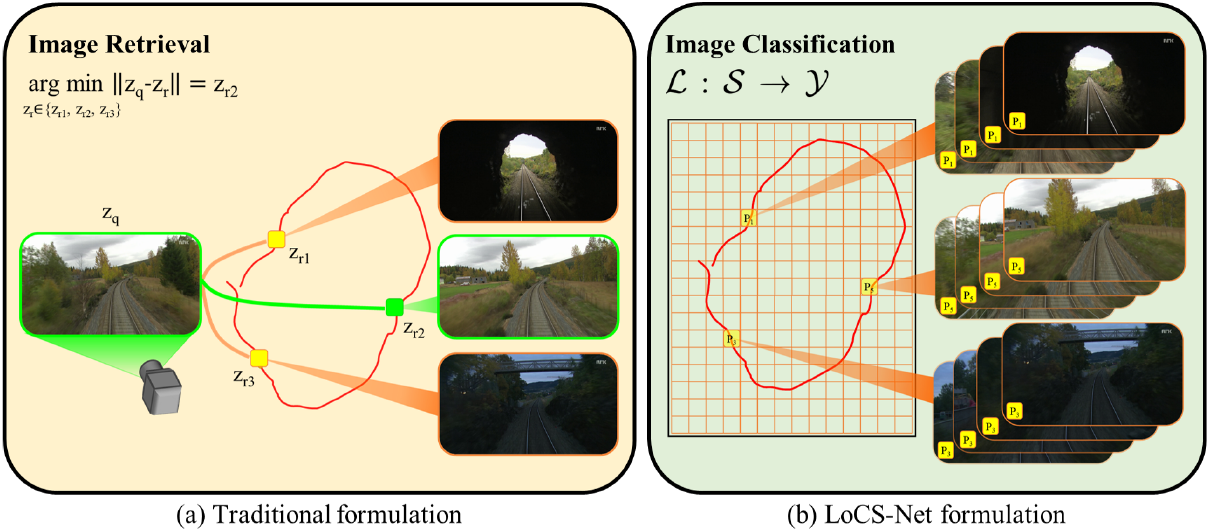
VPR posed as: (a) Image retrieval task, (b) Image classification problem. For both formulations we consider a set of images collected over a trajectory (the red curves) traversing a finite navigation domain. A popular VPR solution is based on generating descriptors for query (z_q_) and reference images (z_r_), which are than compared to each other in terms of a distance (or similarity) metric ∥·∥ in order to retrieve the reference image corresponding to the correct place. For instance, z_r2_ represents the most similar reference image to the given query image. In contrast, the image classification formulation of the VPR task requires an arbitrary discretization of the navigation domain to define the classes *P*_*i*_ (the places where *i ∈* 𝒴 that annotate the images *s ∈* 𝒮. Then, a classifier ℒ is trained to map images *s* ∈ 𝒮to the correct place labels *i ∈* 𝒴.

Consider a set of images, 𝒮 = {*s* ∈ ℝ^*C×H×W*^| *X* (*s*) ∈ 𝒟}, where *C* is the number of color channels, *H* and *W* are the height and width of the images in pixels and 𝒟 is a pre-determined finite horizontal navigation domain. Here, *X* : 𝒮 → *𝒟* is a function that maps the images *s* ∈ 𝒮 to the planar spatial coordinates [*x*_*s*_, *y*_*s*_]^T^ ∈ 𝒟 = *{* [*d*_1_, *d*_2_]^T^ ∈ ℝ^2^ | *x*_min_ ≤ *d*_1_ ≤ *x*_*max*_ ∧ *y*_*min*_≤ *d*_2_ ≤ *y*_*max*_} where *x*_*min*_, *x*_*max*_, *y*_*min*_, and *y*_*max*_ are the bounds of 𝒟 . *X*(*s*) describes the in-plane spatial state of the camera with respect to a local frame of choice when *s ∈* 𝒮 is sampled. The set 𝒴 = {*I* ∈ ℕ | *I* ≤ *N*_*P*_} contains the place labels that annotate *s* ∈ 𝒮 where *N*_*P*_ is the number of assumed places. Each *y* ∈ 𝒴 corresponds to a *P*_*y*_ ⊂ 𝒟 such that *P*_*y*_ ∩ *P*_*i*_ ≡ ∅, *y* ≠ *i* ∧ *i* ∈ 𝒴 . We formulate the VPR task as an image classification problem, where each class is assumed to be mutually exclusive. That is, each image belongs exactly to one class. Our goal is to design a mapping ℒ : 𝒮 → 𝒴 that correctly predicts the place label *y* ∈ 𝒴 of any given *s* ∈ *𝒮* . One should note that the approach we describe here is different than the image retrieval formulation as we want ℒ to predict the place labels instead of directly associating the input images with the reference images.s

### 3.3 Localizing Convolutional Spiking Neural Network

The design of LoCS-Net is defined mainly by two ideas: 1) Discretization of the given finite navigation domain, 2) Leveraging the back-propagation algorithm by adopting the ANN-to-SNN conversion paradigm. We now walk through the details of these ideas together with the architecture of LoCS-Net and its building blocks, LIF neurons.

#### ANN-to-SNN conversion

Due to the discontinuities introduced by discrete spike events, the conventional gradient-descent training techniques need to be modified for spiking neural networks. Various approximation methods have been developed to overcome these discontinuities [61]. One such method is based on the utilization of the rate-based approximations, a.k.a. the tuning curves. Given a loss function, the main idea is to build a network of differentiable rate-based approximation units and solve for the synaptic weights by using an arbitrary version of gradient descent. Once the solution is obtained, the approximation units can be substituted with LIF neurons to use the resulting spiking network during inference as shown in Fig. 3. We utilized NengoDL [62] to implement the aforementioned ANN to SNN conversion methodology. We considered the standard sparse categorical cross entropy as our loss function.

**Figure 3.**
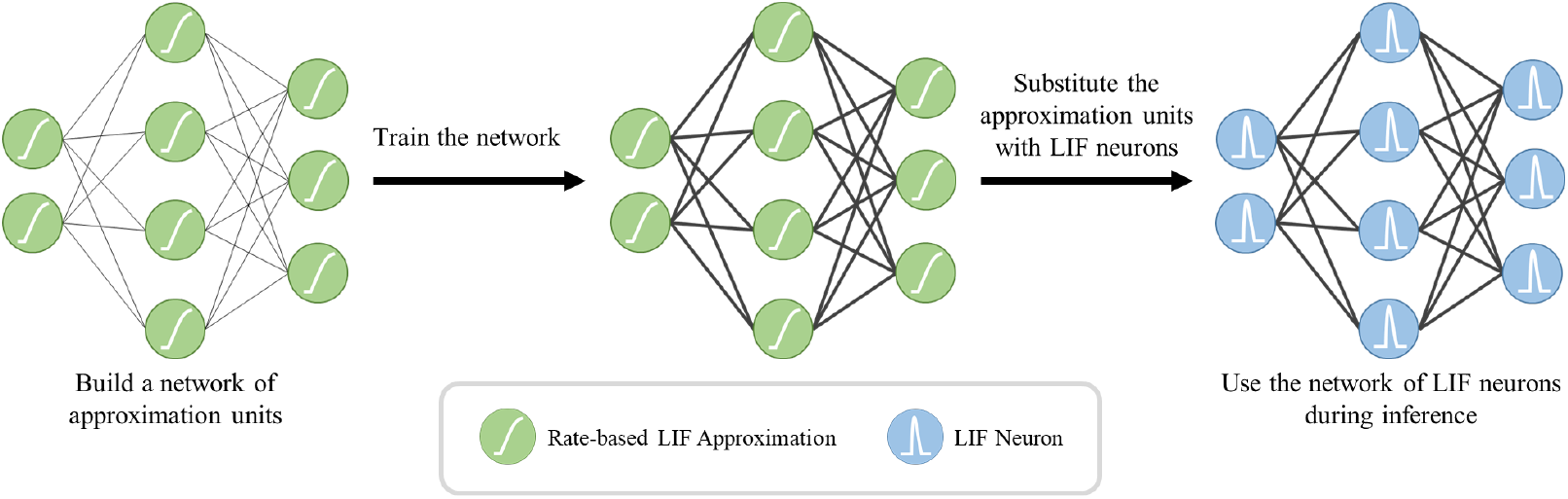
ANN-to-SNN conversion work-flow. We first employ rate-based approximations of the LIF neurons to train our network, since the discontinuous spike event outputs of the original LIF neurons prevents the training of the network through the back-propagation algorithm. After completing the training of this interim network, we substitute the LIF approximations with the original ones while keeping the network topology and the trained weights (bold black lines) the same.

#### LoCS-Net architecture

As depicted in Fig. 1, the LoCS-Net network is composed of a sequence of convolutional layers and a fully connected output layer, in other words the place layer, units of which correspond to distinct places in the environment. We would like to note that LoCSNet consists of 3 convolutional layers and Fig. 1 depicts only the first two convolutions. The number of neurons in the place layer is set to be the number of possible places (*N*_*P*_) as explained in Section 3.2.

## 4 Experiments

### 4.1 Datasets and Evaluation Metrics

We evaluate our proposed approach on the challenging Nordland [22] and Oxford RobotCar (ORC) data [23, 24] following prior work [56]. Although we attempted to follow the data processing directions in [56] for the same training and test data, we couldn’t obtain the same number of places. We obtained 3072 Nordland data [22] places, and 2500 Oxford RobotCar (ORC) data [23, 24] places (set by our grid definition) while considering the complete sun, rain, and dusk traverses used in [56]. In this way, we ensure fairness of the comparison studies. Particularly, we are adopting a conservative evaluation strategy for the ORC data [23, 24] by introducing more places to predict out of a bigger portion of data. Namely, we are considering a more challenging ORC scenario compared to the one in whom the baselines were examined. Appendix A provides the training and the test data specifications yielded by our data pre-processing pipeline. Although our discretization of the ORC domain yields a total of 2500 possible places, the trajectories traversed in that dataset cover a much smaller number of labels. This is because the trajectories were generated by a vehicle traversing the road network, and it was not possible to visit all parts of the spatial domain. There were only 382 unique place labels in the 2014-11-21 16:07 ORC trajectory that we considered for our principal experiments.

We employed standard VPR performance metrics, including the precision-recall curves, area-underthe-precision-recall curves (AUC-PR or AUC) [63, 64], and recall-at-N curves [65, 66] in order to assess the performance of our model.

### 4.2 Experimental Set-Up

We adopt two annotation methods as the Nordland [22] and the ORC data [23, 24] were structured in different ways. Figure 4-A describes the labelling process of the ORC images [23, 24]. As it is shown, we first encapsulated the top-down projection of the path within a rectangular region. Then, we discretize this region to obtain grid tiles, each of which represents a distinct place. These tiles annotate the images sampled within its boundaries. To label the Nordland images [22] we followed the annotation method defined in [56]. As depicted in Figure 4-B, we sample images over a traverse at a pre-determined frequency while each image is corresponding to a unique place.

**Figure 4.**
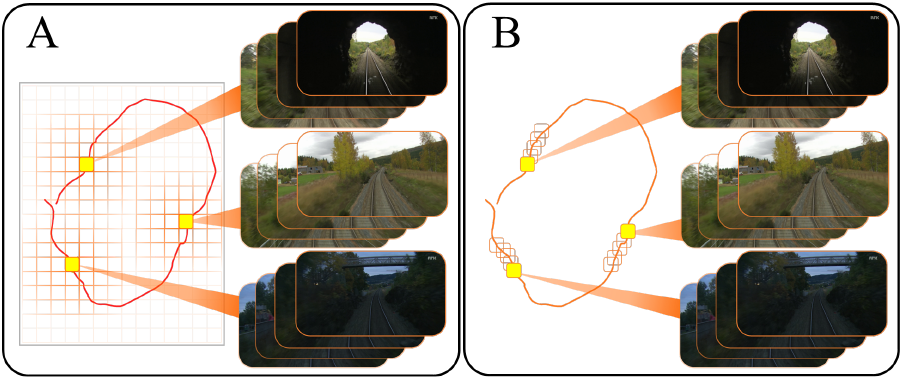
Annotating images. A) Top-down navigation domain is discretized by defining a grid of arbitrary resolution. Each tile of of the grid annotates the images sampled within its boundaries. B) Images are sampled over a traverse at a pre-determined frequency while each image is corresponding to a unique place.

### 4.3 Quantitative Results

We conducted several performance comparisons of LoCS-Net with state-of-the-art conventional and SNN-based VPR methods. We compared our method with Sum-of-Absolute-Differences (SAD) [67], NetVLAD [48], the current best-performing SNN method, Ensemble SNNs [56], and a previous SNN method by the same authors, Weighted Assignment SNN (WNA) [55]. This selection of methods was determined by following the choice of baselines considered by the current SOTA SNN-based VPR work [56].

Figure 5 and Table 1 summarize the VPR performance of LoCS-Net along with the reference methods. We observed that LoCS-Net outperformed the SNN-based method WNA [55] on both the Nordland [22] and ORC dataset [23, 24] by a large margin (78.2% and 34.8% respectively) in terms of precision at 100% recall. LoCS-Net also outperformed Ensemble SNNs [56] by more than 25 percentage points on the Nordland dataset [22] while being worse by 6 percentage points on the ORC data [23, 24] in terms of precision at 100% recall as reported in Table 1 of [56]. We believe that the gradient mismatch between the LIF neurons [21] and their rate approximations [20] has a significant contribution to the reduced performance of LoCS-Net in this case, as also mentioned by [68]. LoCS-Net outperformed both methods on both datasets in terms of area-under-the-precision-recall curves, while taking significantly less inference time as shown in Tab. 1. For fair comparison, we followed the preprocessing steps as described in [56]. However, we were not able to independently reproduce the results of the methods reported in [56]. Therefore, we extracted the results from Figures 2 and 3 of [56]. Moreover, we used the benchmark tool created by Zaffer et al. [69] in order to compute the MIT values of NetVLAD [48].

**Table 1:** VPR performance comparison in terms Precision at 100% Recall (P@100%R), area-under-the-precision-recall curves (AUC), and mean inference time (MIT).

| Method | Approach | Nordland |  |  | ORC |  |  |
| --- | --- | --- | --- | --- | --- | --- | --- |
| | | $P@100\%R$ | AUC | MIT[ms] | $P@100\%R$ | AUC | MIT[ms] |
| LoCS-Net (ours) | SNN | **78.2%** | **0.967** | **171** | 34.8% | **0.616** | **193** |
| WNA [55] | SNN | 0.3% | 0.002 | 564 | 4.0% | 0.035 | 564 |
| Ensemble SNNs [56] | SNN | 52.6% | 0.753 | 379 | **40.5%** | 0.613 | 379 |
| SAD [67] | Heuristics | 45.1% | 0.668 | - | 41.3% | 0.594 | - |
| NetVLAD [48] | ANN | 35.1% | 0.425 | 958 | 44.8% | 0.659 | 1087 |

**Figure 5.**
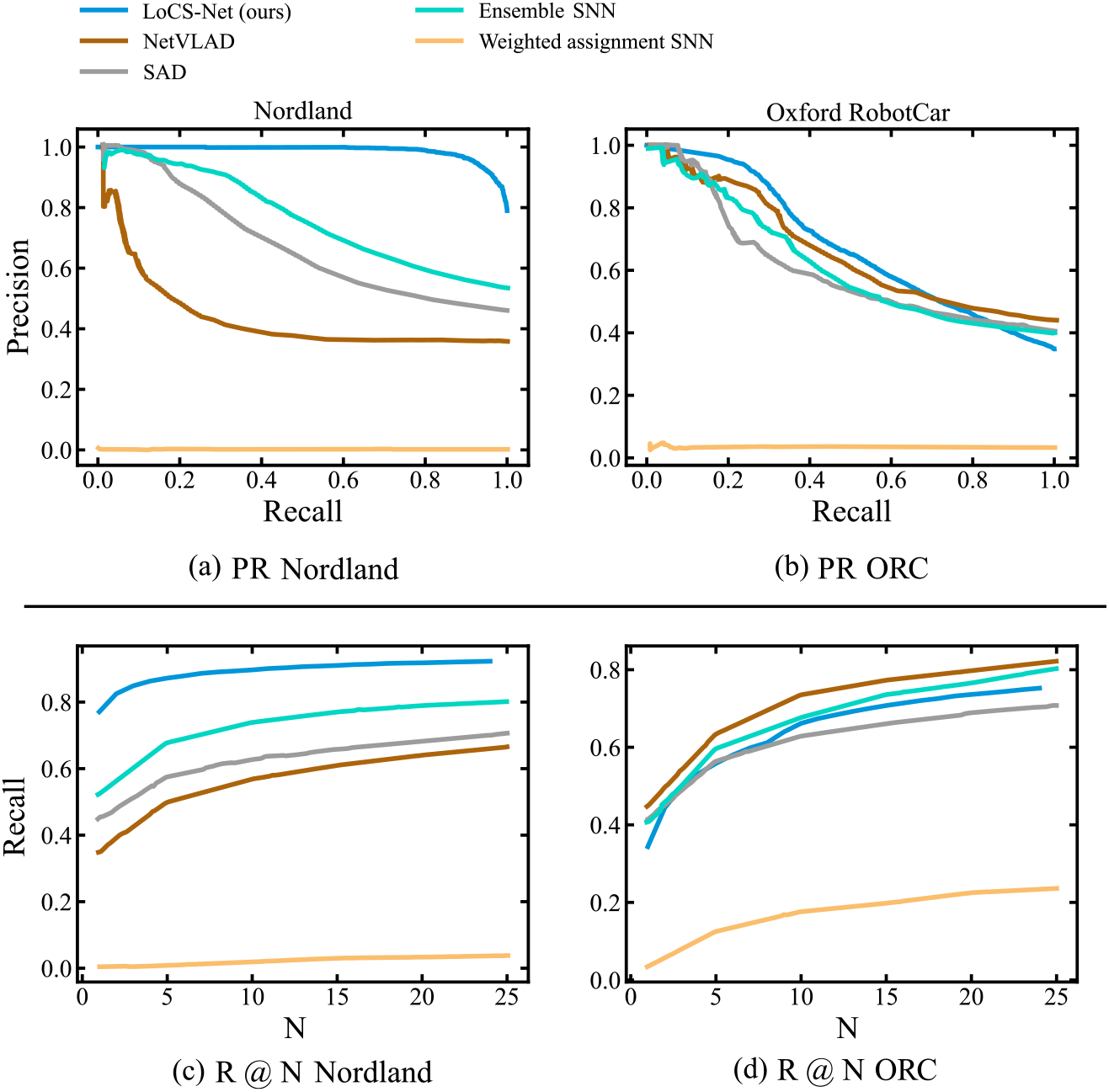
Precision-Recall and Recall @ N curves for the baseline VPR methods and LoCS-Net. (a) and (c): Nordland data [22], (b) and (d): ORC Data [23, 24].

LoCS-Net also surpassed the performance of the heuristics-based SAD [67] by more than 30 percentage points on the Nordland dataset [22], while being slightly worse (6 percentage points) on the ORC dataset [23, 24]. Our approach outperformed the ANN-based NetVLAD [48] by more than 40 percentage points on the Nordland dataset [22], while performing worse on the ORC dataset [23, 24] by 10 percentage points. We observe a consistent poor performance of our method over the ORC dataset [23, 24]. In addition to the gradient mismatch we mentioned above, we hypothesize that the more dynamic scene content of the ORC images [23, 24] impedes better VPR performance of LoCS-Net. We would also like to emphasize that NetVLAD [48] experiments were conducted with input images of 640x360 pixels, which might be more advantageous as there would be less information loss. However, Table 1 highlights the fastest inference time (MIT) of LoCS-Net among all of its competitors. We note that the particular selection of the ORC trajectories [23, 24] was especially challenging, as there was a large variance in lighting and road conditions.

Furthermore, Figure 5 presents the Recall @ N curves obtained from the evaluations of the methods over the Nordland [22] and ORC datasets [23, 24]. LoCS-Net consistently yields the best Recall @ N performance compared with both conventional and SNN method on the Nordland data [22], and has a Recall @ N performance similar to the other methods on the ORC dataset [23, 24]. These results indicate good scalability of the LoCS-Net model over thousands of locations, while the inference is computationally efficient, as illustrated by Table 1.

Finally, in order to understand how well the proposed method would generalize to other datasets, as well as different input image sizes, we investigated the performance of LoCS-Net over a broad range of standard VPR benchmark data sets, as well as with different input image sizes. The results of these experiments confirmed that LoCS-Net performs at the top of its class on a number of standard benchmark datasets. Detailed results are presented in Appendix C.

## 5 Conclusion

In this work, we formulate visual place recognition as a classification problem and develop LoCS-Net, a convolutional SNN to solve VPR tasks with challenging real-world datasets. Our approach leverages ANN-to-SNN conversion and back-propagation for tractable training, by using rate-based approximations of leaky integrate-and-fire (LIF) neurons. The proposed method outperforms the state-of-the-art ANNs and SNNs on the challenging Nordland dataset [22] and achieves competitive performance on the ORC dataset [23, 24], while being easier to train and faster in both training and inference as compared to other ANN and SNN-based methods. The current work exhibits a significant potential towards the deployment of SNN-based VPR systems on robotics platforms for real-time localization.

### Limitations

We would like to emphasize that this manuscript proposes LoCS-Net as a environment-specific VPR solution. In that sense, providing a long-term general spatial memory as a global VPR solution is beyond the capabilities of the current work. In addition, LoCS-Net’s performance is sensitive to the definition of places, which in turn may require the implementation of domain-specific discretization techniques to maximize the performance of LoCS-Net over different navigation environments. As discussed in Section 4.3, LoCS-Net doesn’t perform as well over the Oxford RobotCar [23, 24] data as compared with the Nordland [22] dataset. This might be due to the LIF neuron approximation errors as well as the significantly varying lighting and road conditions of the ORC traverses [23, 24], which suggests the lack of robustness to such dynamic scenes.

## A Pre-processed Data Specifications

Our pre-processing of Nordland [22] and ORC [23, 24] datasets produced the training and the test data with specifications listed in Table 2. The pre-processing code will be available upon the request of the readers.

**Table 2:** LoCS-Net training and test data specifications.

| Specifications \ Dataset | Nordland | Oxford RobotCar (ORC) |
| --- | --- | --- |
| Train size [# of images] | 6144 | 72207 |
| Test size [# of images] | 3072 | 18982 |
| # of labels | 3072 | 2500 |
| # of unique labels | 3072 | 382 |

## B The Governing Equations of the LIF Neurons

### LIF model

Unlike standard artificial neurons, which are defined by time-independent differentiable non-linear transfer functions with continuous outputs, spiking neurons have time-dependent dynamics that aim to capture the information processing in the biological neural systems by emitting discrete pulses[21]. Equation (1) describes the dynamics of a LIF neuron.

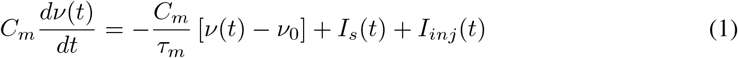

where *C*_*m*_ is the membrane capacitance, *τ*_*m*_ is the passive membrane time constant, and *ν*_0_ is the resting potential. Above formulation considers a resetting scalar state variable, the membrane potential *ν*(*t*), which will be reinitialized at *ν*(*t*) = *ν*_*reset*_ after reaching a threshold, *ν*(*t*) = *ν*_*th*_.

Whenever the re-initialization happens at time *t* = *t*_*spike*_, the output of the LIF neuron (*o*(*t*)) will be an impulse signal of unity. We name this a spike event. One can express a spike event of a LIF neuron by Equation (2), which incorporates Dirac’s delta function centered at the time of re-initialization.

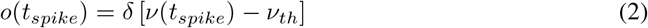

The right hand side of Equation (1) includes three terms: 1) An exponential decay term (a.k.a the passive membrane leak), 2) *I*_*s*_(*t*), the sum of incoming synaptic currents, which are mostly unit impulses filtered through a first order delay and/or multiplied by some scalar, and finally 3) An injection term, *I*_*inj*_(*t*), that describes the input currents other than synaptic currents. This can be some bias representing the background noise in the corresponding neural system, or just some external input.

### Rate-based approximations of the LIF neurons

Solving the sub-threshold dynamics described by Equation (1) for the firing rate, *ρ*[*I*_*s*_(*t*)], of a LIF neuron and assuming *I*_*inj*_(*t*) = 0 for all *t ≥* 0 yields the following.

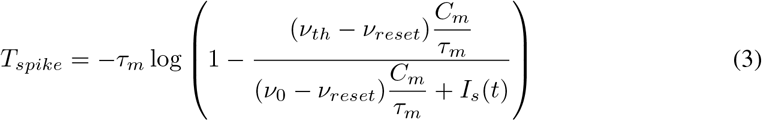

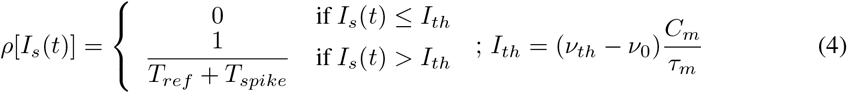

*T*_*ref*_ is the refractory period, which is the time it takes a neuron to start accepting input currents after a spike event. *T*_*spike*_ is the time it takes a neuron to reach *ν*_*th*_ from *ν*_*reset*_ after a spike event at some *t* = *t*^*′*^ given *ν*_*th*_ *< I*_*s*_(*t*) = *c* ∈ ℝ, *t*^*′*^ *< t ≤ t*^*′*^ + *T*_*spike*_. Equations (3) and (4) describe the response curve of a LIF neuron, which has a discontinuous and unbounded derivative (*∂ρ/∂I*_*s*_) at *I*_*s*_ = (*ν*_*th*_ *− ν*_0_)*C*_*m*_*/τ*_*m*_. However, one can modify Equation (4) as described by [20] in order to obtain a smooth rate-based LIF approximation.

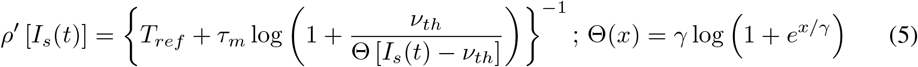

where *γ* is the smoothing factor of choice.

## C Additional Experiments

In order to understand how much the proposed method would generalize to other datasets, we investigated the performance of LoCS-Net over a broad range of standard VPR benchmark datasets, as well as with different input image sizes. We considered a wide variety of baselines other than our main SNN competitors [55, 56].

We utilized a standardized benchmarking platform for visual place recognition (VPR) systems (VPR-Bench [69]) to evaluate our method at different input resolutions as well as conduct comparisons with other baselines [9, 48, 70, 71, 72, 73, 8, 7, 41, 74] over additional datasets [75, 76, 77, 78]. We didn’t consider datasets where input images are assigned multiple labels as such annotation is not compatible with our VPR formulation. We present the VPR performance of LoCS-Net and the other baselines over the aforementioned datasets in Fig. 6 and Table 3. Fig. 6 demonstrates the precision-recall (PR) curves of the 3 best performing baselines for easier read of the figure. Table 3 covers all of the baselines’ VPR performance over the additional datasets in terms of Precision at 100% Recall (P@100%R) values. We would like to note that we didn’t implement any of the baselines, but we incorporated the results reported in [69]. Fig. 6 and Table 3 highlight the superior VPR performance of LoCS-Net and/or its variants over a variety of indoor and outdoor datasets included in VPR-Bench. We initially used only the Oxford Robot Car and Nordland datasets as our primary goal was to maintain a fair comparison with the current state-of-the-art (SOTA) spiking neural networks (SNNs) [56, 55]. In that sense we closely followed the experimental setups in [56, 55].

**Table 3:** VPR performance comparison over the additional datasets included in VPR-Bench [69]. Here we include the complete list of Precision at 100% Recall (P@100%R) values obtained from the experiments.

| Method | Approach | Input | P@100%R |  |  |  |
| --- | --- | --- | --- | --- | --- | --- |
|  |  |  | Synthia[78] | Living[77] | Gardens Point[76] | Corridor[75] |
| LoCS-Net (ours) | SNN | 28x28 | 0.972 | 0.966 | 0.997 | 0.983 |
| LoCS-Net (ours) | SNN | 56x56 | 0.965 | **1.000** | **1.000** | **1.000** |
| LoCS-Net (ours) | SNN | 112x112 | **0.980** | **1.000** | **1.000** | **1.000** |
| NetVLAD [48] | ANN | 640x480 | 0.865 | 0.938 | 0.582 | 0.676 |
| RegionVLAD [70] | ANN | 227x227 | 0.621 | 0.690 | 0.432 | 0.436 |
| DenseVLAD [74] | View Synthesis | 640x480 | 0.911 | 0.968 | 0.516 | 0.716 |
| AlexNet-VPR [9] | ANN | 227x227 | 0.622 | 0.613 | 0.272 | 0.303 |
| HOG [72] | Heuristics | 512x512 | 0.976 | **1.000** | 0.197 | 0.5 |
| CoHOG [71] | Heuristics | 512x512 | 0.773 | 0.844 | 0.390 | 0.622 |
| AMOSNet [8] | ANN | 227x227 | 0.844 | 0.750 | 0.477 | 0.887 |
| HybridNet [8] | ANN | 227x227 | 0.889 | 0.714 | 0.457 | 0.901 |
| CALC [73] | ANN | 120x160 | 0.685 | 0.520 | 0.206 | 0.462 |
| AP-GeM-r101 [41] | ANN | 640x480 | 0.865 | 0.906 | 0.530 | 0.536 |

**Figure 6.**
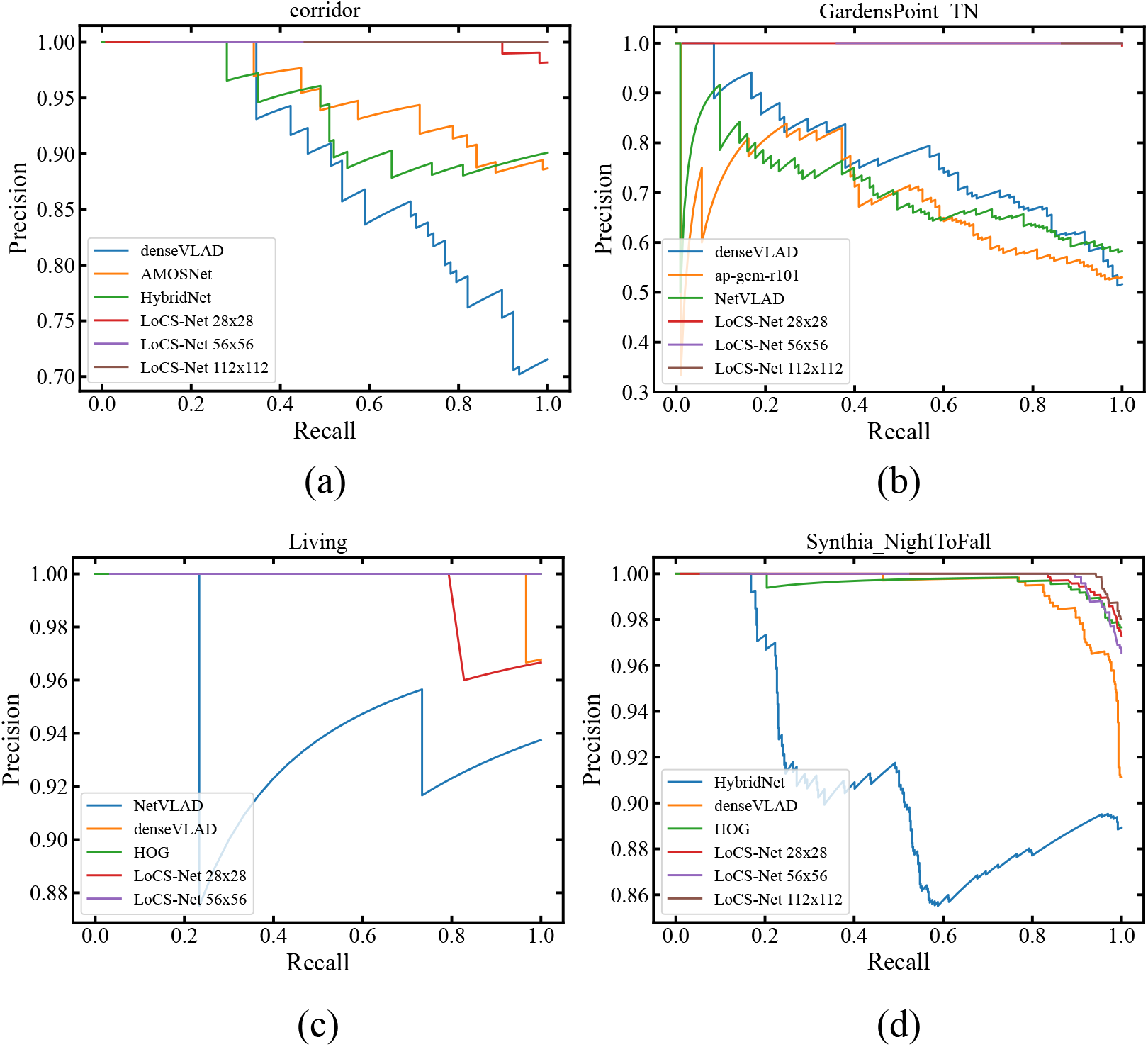
Precision-Recall (PR) curve comparisons of LoCS-Net and other baselines over additional datasets included in [69]. We compared LoCS-Net and its different input resolution variants’ PR curves to a number of different baselines’ PR curves over 4 additional datasets: (a) Corridor [75], (b) Gardens Point [76], (c) Living-room [77], (d) Synthia [78]. For better readability of the plots, we included the PR curves of 3 best performing baselines. LoCS-Net demonstrates the best VPR performance over all of the datasets considered here.

To assess the generalizability of our model, we tested LoCS-Net on additional ORC trajectories [23, 24]. We chose 2015-02-20 16:34 so as to be maximally close to the season and time of the day of the original 2014-11-21 16:07 dataset used by closely related SNN-based models [55, 56]. Fig. 7 dmonstrates improved performance of LoCS-Net (from 34.8% to over 50% *P* @100%*R*) over the 2015-02-20 16:34 trajectory compared to the 2014-11-21 16:07 one.

**Figure 7.**
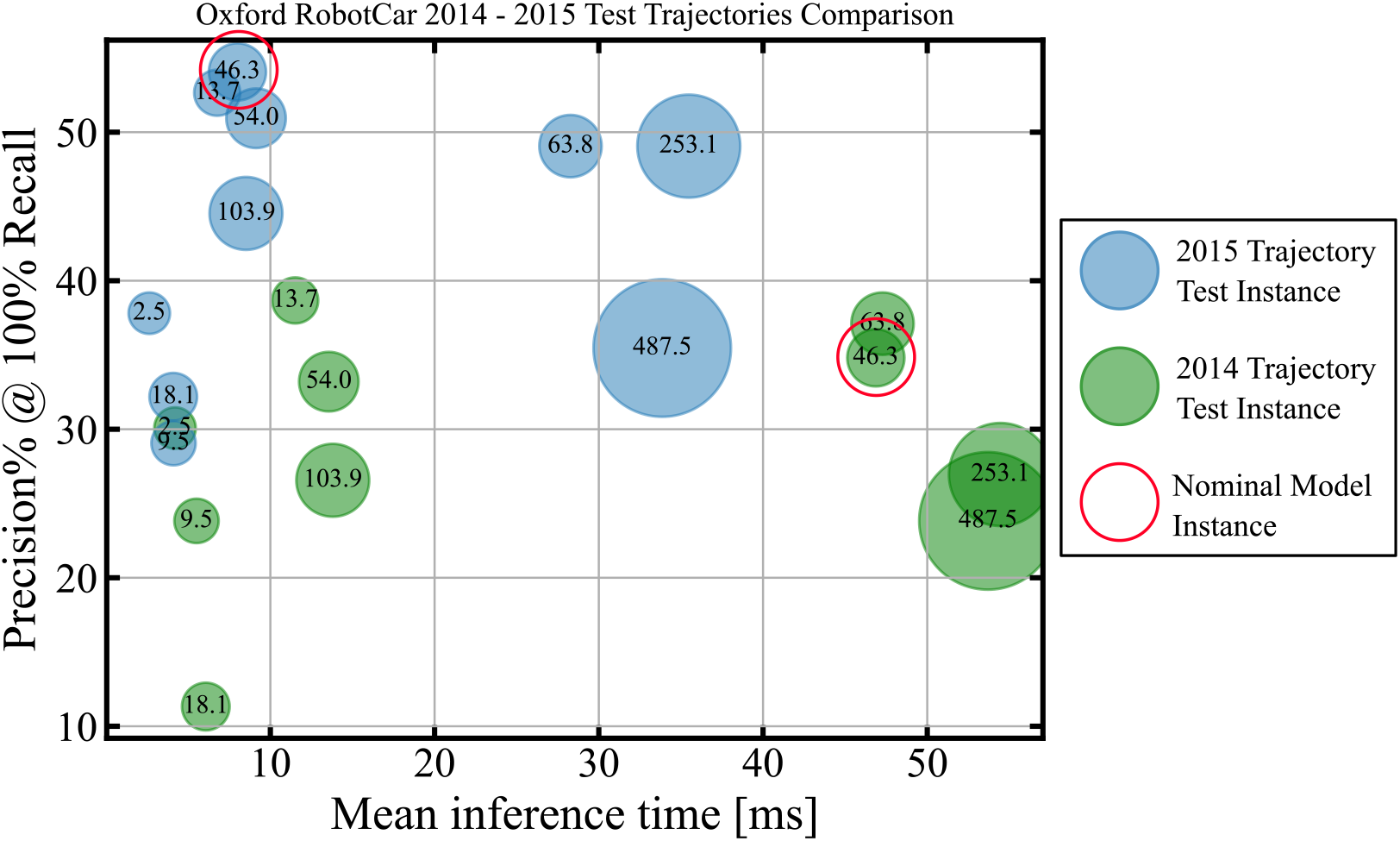
Precision at 100% Recall and Mean Inference Time (MIT) achieved by LoCS-Net variants over Oxford RobotCar [23, 24] 2014-11-21 16:07 and 2015-02-20 16:34 trajectories with different input and output resolutions. Each green and blue circle correspond to an instance of LoCS-Net variant tested over 2014-11-21 16:07 and 2015-02-20 16:34 trajectories, respectively. Numbers in the circles stand for the total number of trainable parameters in millions whereas the circle sizes get bigger with the increasing number of trainable parameters. Red circle indicates the nominal LoCS-Net model of 56x56 input resolution and 50x50 grid resolution. Results suggest that better VPR perfomance can be achieved with smaller networks.

Furthermore, we conducted additional experiments spanning a range of different input and grid resolutions, results of which are demonstrated in Figs. 7 and 8. In Figs. 7 and 8 the model sizes are given in terms of number of trainable parameters in millions. Tables 4 and 5 list the input resolutions, output grid resolutions, and the Nordland data sampling rate (the skip parameter) that correspond to the numbers of trainable parameters. The results confirm the that LoCS-Net provides SOTA SNN performance for the VPR task. LoCS-Net yielded improved performance (compared to the nominal model) over both the Nordland data [22] and the ORC data [23, 24] for some combinations of input-output resolution. In fact, a smaller model of 28.6 million trainable parameters exhibits the best performance over the Nordland data as shown in Fig. 8. Varying the input and grid resolutions did result in a range of different MITs that are reported in Figs. 7 and 8. Figs. 7 and 8 show mostly an increasing MIT by increasing model size with some counter-intuitive instances. For example, in Fig. 8 shows that the model with 532 million parameters has smaller MIT than the one with 133.7 million parameters. We carried out the numerical experiments on a shared cluster. Thus, it is possible that variance in the system load had affected the MIT values.

**Table 4:** The input and grid resolutions that correspond to the numbers of trainable parameters of LoCS-Net variants tested over ORC 2014-11-21 16:07 and 2015-02-20 16:34 trajectories. The number of parameters are given in millions. The entry with bold font points the nominal model that is used for generating the results reported in the submitted manuscript.

| Input Resolution \ Grid Resolution | 27x27 | 50x50 | 54x54 | 75x75 |
| --- | --- | --- | --- | --- |
| 28 | 2.5 M | - | 9.5 | 18.1 |
| 56 | 13.7 M | **46.3 M** | 54 | 103.9 |
| 112 | 63.8 M | - | 253.1 | 487.5 |

**Table 5:** The input resolution and the skip parameters that correspond to the numbers of trainable parameters of LoCS-Net variants tested over the Nordland data [22]. The number of parameters are given in millions. The entry with bold font points the nominal model that is used for generating the results reported in the submitted manuscript. The skip parameter defines the sampling rate of the data. Say the skip rate is N, then every N^*th*^ image will be collected. In Nordland data, every image is considered as a place. Thus, after sampling of the entire data, the resulting number of images would be also the number of LoCS-Net’s output layer neurons.

| Input Resolution \ Skip Parameter | 4 | 8 | 16 |
| --- | --- | --- | --- |
| 28 | 19.8 M | 10.0 M | 5.0 M |
| 56 | 113.5 M | **56.9 M** | 28.6 M |
| 112 | 532.3 M | 266.6 M | 113.7 M |
| 224 | - | - | 576.1 M |

**Figure 8.**
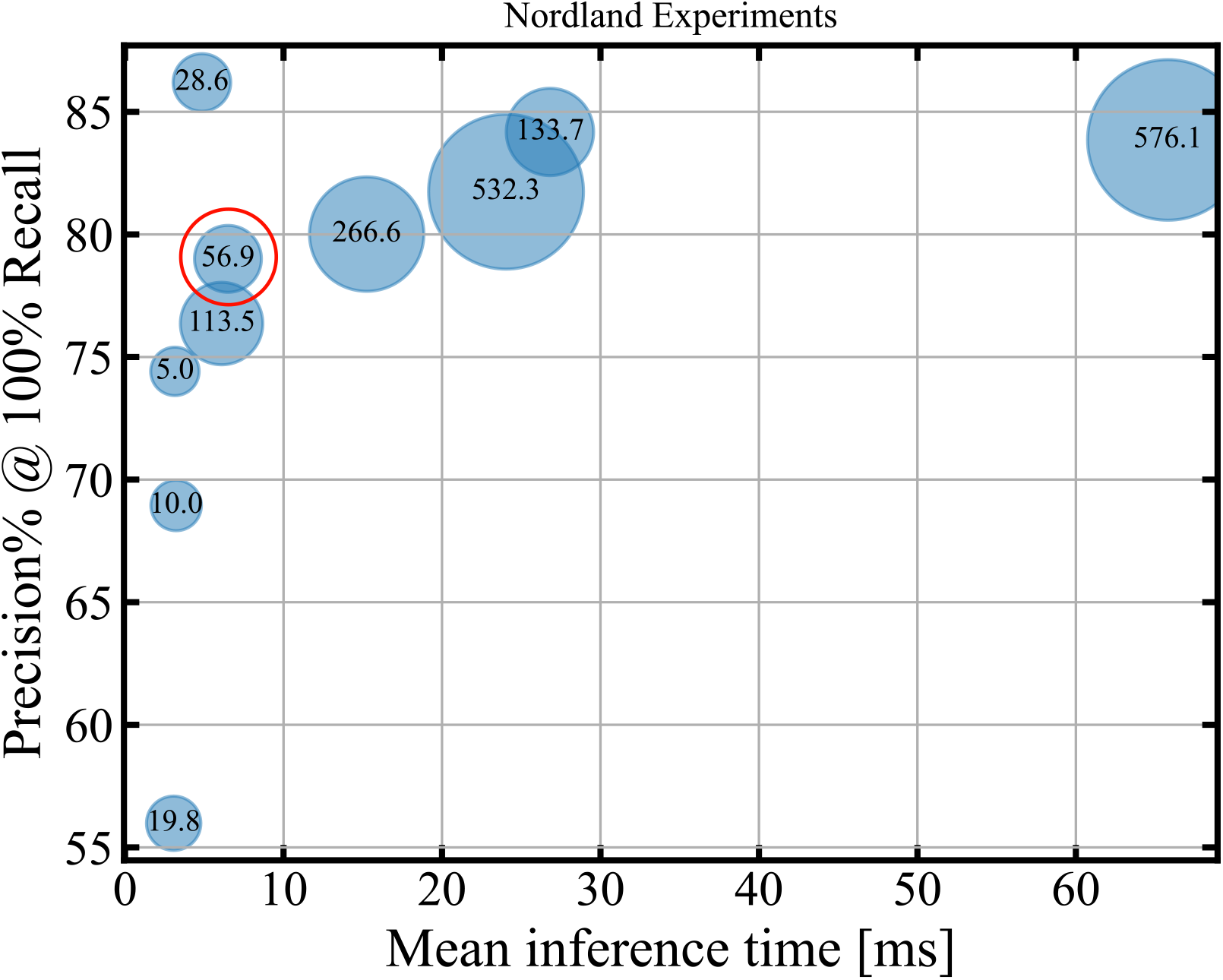
Precision at 100% Recall and Mean Inference Time (MIT) achieved by LoCS-Net variants over the Nordland dataset [22]. Each blue circle corresponds to an instance of LoCS-Net variant tested over the Nordland dataset. Numbers in the circles stand for the total number of trainable parameters in millions whereas the circle size get bigger with the increasing number of trainable parameters. Red circle indicates the nominal LoCS-Net model of 56x56 input resolution and skip parameter 8. Results depict that better VPR perfomance can be achieved with smaller networks.

We further sought to understand the distribution of LoCS-Net’s prediction errors on the ORC dataset. We quantify the prediction error in terms of Manhattan Distance, as illustrated in Fig. 9. Most of the erroneous place predictions were within 1-Manhattan Distance, which indicates that most of the model’s false predictions are relatively close to the ground truth place labels. We did not perform the same analysis for Nordland data as it doesn’t utilize a grid-based labelling structure as the Oxford RobotCar data.

**Figure 9.**
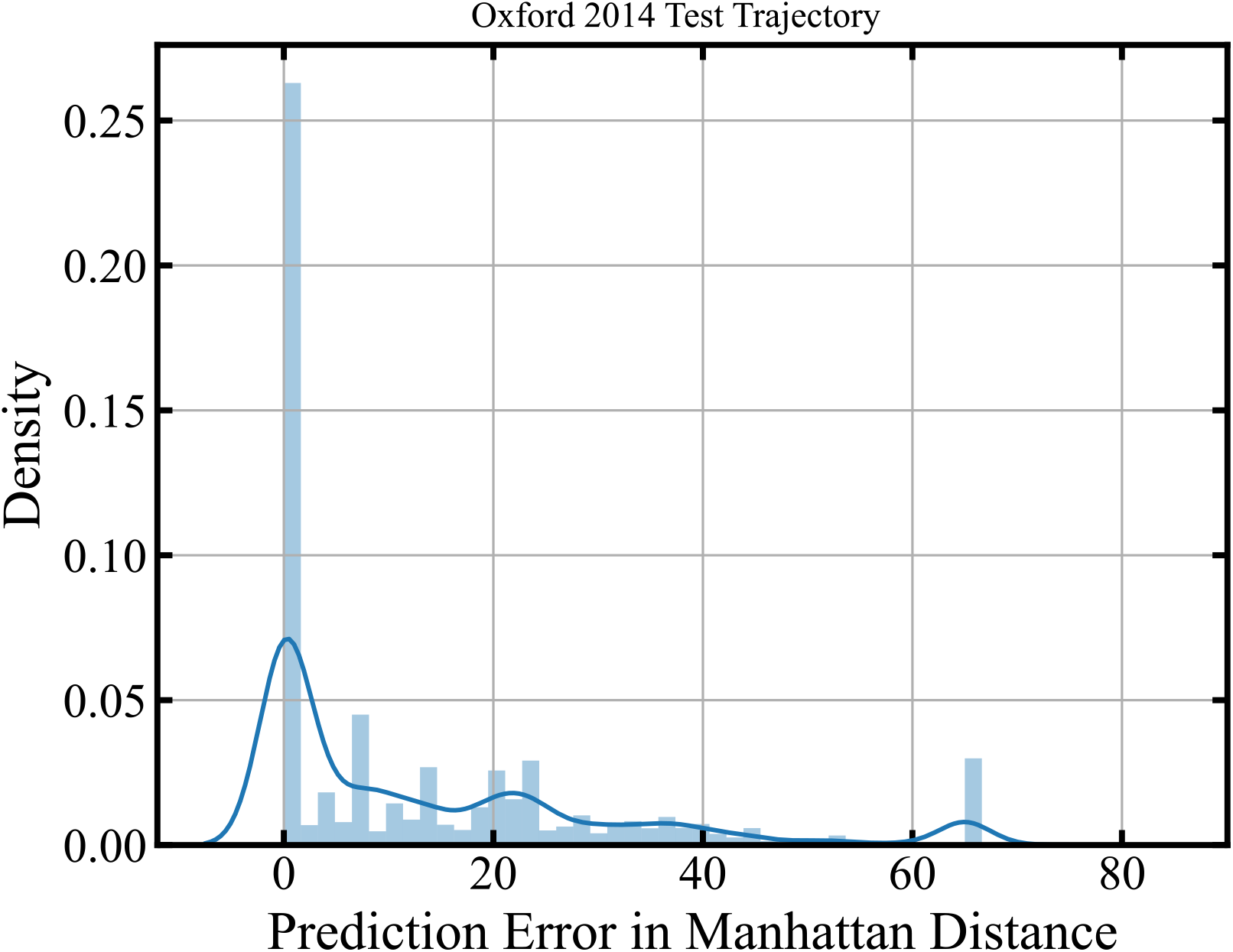
Prediction error distribution of LoCS-Net over the ORC dataset. Most of the erro-neous place predictions of LoCS-Net are within 1-Manhattan Distance, which implies that most of the model’s false predictions are relatively close to the ground truth place labels.

In addition to the above empirical analysis, we carried out a number of experiments comparing LoCS-Net to a traditional convolutional neural network (CNN) of rectified linear units (ReLUs) with the same architecture. We evaluated the conventional CNN over the same data with LoCS-Net. Table 6 shows the corresponding results. As reported in Table 6, the performance of the conventional CNN is surpassed by LoCS-Net over the ORC trajectories while demonstrating similar VPR performance over the Nordland data. The difference in performance suggests that there are more factors than just the multi-class classification formulation.

**Table 6:**
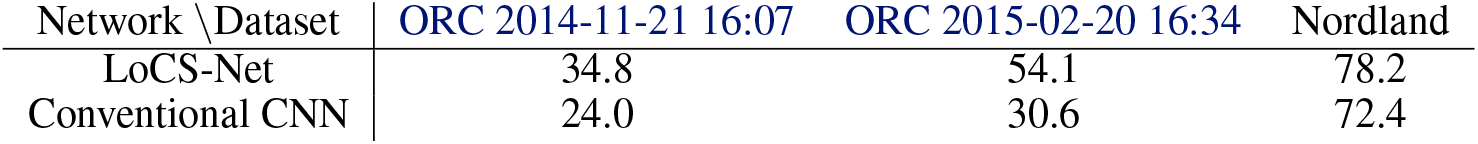
Comparing LoCS-Net to its conventional CNN equivalent with respect to Precision% @ 100% Recall. The traditional CNN has the same architecture with LoCS-Net, except it is of ReLU activations.

